# Soybean and cotton spermosphere soil microbiome shows dominance of soil-borne copiotrophs

**DOI:** 10.1101/2023.01.23.525219

**Authors:** Oluwakemisola E. Olofintila, Zachary A. Noel

## Abstract

The spermosphere is the transient, immediate zone of soil around imbibing and germinating seeds. It represents a habitat where there is contact between seed-associated microbes and soil microbes, but is studied less compared to other plant habitats. Previous studies on spermosphere microbiology were primarily culture-based or did not sample the spermosphere soil as initially defined in space and time. Thus, the objectives of this study were to develop an efficient strategy to collect spermosphere soils around imbibing soybean and cotton in non-sterile soil and investigate changes in microbial communities. The method employed sufficiently collected spermosphere soil as initially defined in space by constraining the soil sampled with a cork borer and confining the soil to a 12-well microtiter plate. Spermosphere prokaryote composition changed over time and depended on the crop within six hours after seeds were sown. By 12 to 18 hours, crops had unique microbial communities in spermosphere soils. Prokaryote evenness dropped following seed imbibition with the proliferation of copiotrophic soil bacteria. Due to their long history of plant growth promotion, prokaryote OTUs in *Bacillus, Paenibacillus, Burkholderia, Massilia, Azospirillum*, and *Pseudomonas* were notable genera enriched. Fungi and prokaryotes were hub taxa in cotton and soybean spermosphere networks. Additionally, the enriched taxa were not hubs in networks, suggesting other taxa besides those enriched may be important for spermosphere communities. Overall, this study advances knowledge in the assembly of the plant microbiome early in a plant’s life, which may have plant health implications in more mature plant growth stages.

## Introduction

When a seed is sown, it imbibes water and releases nutrient-rich seed exudates that fuel interactions between soil and seed-associated microbes in a plant habitat called the spermosphere (Nelson 2004; Nelson 2018; Schiltz et al. 2015; Shade et al. 2017; Windstam and Nelson 2008). Seed exudates have long been recognized to stimulate microbial growth, including a direct link to facilitating pathogen growth chemotactically towards seeds (Nelson 2004; Short 1976; Slykhuis 1947; Stanghellini 1971). The spermosphere or “spermatosphere” was first described by Slykhuis (1947), who observed the inhibition of a fungal pathogen by three fungal species around a germinating seed. Verona (1958) defined the same habitat as a “zone of elevated microbial activity” around a germinating seed. Nelson (Nelson 1986, 1988; 2004; van Dijk and Nelson 2000; Windstam and Nelson 2008) more formally defined the spermosphere as “the short-lived, rapidly-changing and microbiologically dynamic five-to-ten-millimeter zone of soil around a germinating seed”, which is the definition we adhere to in this study. Despite its importance for plant health outcomes, the spermosphere is less studied than other plant-associated habitats such as the rhizosphere or phyllosphere (Aziz et al. 2022; Schiltz et al. 2015; Shade et al. 2017).

Seed germination occurs in three distinct phases. Phase I is a physical process characterized by seed imbibition and fast carbon-rich exudate release into the soil hours after seeds are sown. The highest levels of exudate release are completed in as little as six hours (Lynch 1978; Nelson 2004; Simon and Raja Harun 1972). The initial phase of exudate release is followed by a plateau characterizing phase II, then radical emergence, which begins the formation of the rhizosphere, and more exudate release in phase III (Nelson 2004; Schiltz et al. 2015).

The spermosphere represents a critical zone for establishing vertically inherited seed microbes, and horizontal interactions between soil and seed-associated microbes (Chesneau et al. 2022; Chesneau et al. 2020; Rochefort et al. 2021; Shade et al. 2017; Simonin et al. 2022). The outcome of these interactions can affect the life or death of the plant soon after sowing seeds (McKellar and Nelson 2003; Windstam and Nelson 2008). For example, soil-borne *Pythium* can fully colonize and kill germinating seeds of various crop species within 12-24 hours (Hayman 1969; Nelson 1986, 1988; Stanghellini 1973).

Spermosphere pathogens still cause millions of dollars in crop loss yearly (Allen et al. 2017; Bradley et al. 2021; Mueller et al. 2020). Because of enhanced genetics and other factors, seed ranks first or second in operating costs borne by soybean and cotton farmers each year (USDA-ERS 2022). Additionally, trends toward earlier planting dates, increased frequency of heavy pulsed rain events, and variable temperature conditions experienced by farmers at planting can create soil moisture and temperature conditions that stress the germinating seed (Morris et al. 2021; Munkvold 1999; Roth et al. 2020). Conservation tillage (low or no-till) can also lead to harboring plant pathogens on plant debris left in the field from the previous growing season (Bockus and Shroyer 1998). Consequently, the protection of seeds from pathogens that specialize in spermosphere colonization is vital to improved crop productivity.

Seed and seedling pathogens are primarily managed with chemical seed coatings containing fungicides and oomicides. However, improved knowledge on spermosphere microbiology and ecology would support the successful inclusion of alternative strategies to chemical seed treatments. For example, biocontrol of *Pythium* from seed-applied *Enterobacter cloaceae* could be achieved by metabolizing long-chain fatty acids, which otherwise stimulated the germination of *Pythium* sporangium (Kageyama and Nelson 2003; van Dijk and Nelson 2000; Windstam and Nelson 2008). Studies on the spermosphere have either been culture-based or have more recently focused on the contribution of the indigenous seed microbiome by using sterile or soilless growth conditions or pre-imbibed or pre-germinated seeds, which do not sample the spatial and temporal properties of the spermosphere soil (Barret et al. 2015; Johnston-Monje et al. 2016; Moroenyane et al. 2021; Rochefort et al. 2021). Indeed, natural seed-associated epiphytes and endophytes compete with pathogens (Smith et al. 1999; Torres-Cortes et al. 2019). While commendable, these studies largely ignore the influence of the initial seed exudate release on the spermosphere soil microbes. Therefore, a mechanistic understanding of the complex interactions in spermosphere soil will aid in novel treatments for seed and seedling pathogens and help our understanding of plant microbiome assembly.

However, one major challenge in studying spermosphere soil using high-throughput culture-independent techniques may be a lack of a quick and efficient method of collecting spermosphere soil (Schiltz et al. 2015). Here, we aimed to capture changes in microbial diversity in the spermosphere soils as soybean and cotton seeds underwent phases I and II of seed germination (i.e., pre radical emergence). We sampled the spermosphere soil of cotton and soybean by constraining the soil zone within wells of a 12-well plate and sampling precisely three to six millimeters of soil around an imbibing seed with an appropriately sized cork borer, extracted DNA, and sequenced the 16S and ITS from cotton and soybean spermosphere soil. We hypothesized that seeds would imbibe water rapidly and follow previously established phases of exudate release, which would alter microbial diversity and co-occurrence patterns. We also hypothesized that spermosphere soils would be distinct based on crop species. Therefore, the objectives of this study were twofold: 1) characterize the bacterial and fungal microbial communities associated with cotton and soybean spermosphere soil compared to control soil and 2) determine how microbial diversity and co-occurrences change over time as a seed imbibes water.

## Materials and Methods

### Soil collection and preparation

The soil used in this study was collected from a field used for cotton, soybean, and corn rotation from Prattville Agricultural Research Unit in Prattville, Alabama (32.42533, -86.4452), since this soil showed consistent emergence of both cotton and soybean in preliminary experiments (*data not shown*), and was not known to contain a high abundance of any specific seedling pathogen. Approximately three liters of soil from the top ten centimeters were collected and transported to the lab. The soil was sieved to eliminate stones and pebbles and air-dried for 24 hours to ensure homogeneity in water content. The soil was used immediately after air drying. Six to seven grams (6 ml) of soil was transferred to each well of the 12-well microtiter plates (VWR American cat no.:10861-556, USA), containing three one-millimeter holes in the bottom of all wells for drainage. Each well in the 12-well microtiter plates measured a total volume of 6.8 ml, depth of each well of a 12-well plate was 15 mm with a width of 23 mm. Each well containing soil was watered with 1.5 ml of sterile water (25% soil moisture), and the water was allowed to circulate for one hour before the seeds were sown.

Nontreated Williams-82 soybean or nontreated delinted Delta Pine 1646 B2XF cotton were used in this study and were sorted to discard discolored seeds or seeds with cracked seed coats (Nelson 1986). The weight of individual dry seeds was recorded before use and after imbibition to record how much water was imbibed. The initial weight of soybean seeds was between 170 and 250 mg, and cotton seeds weighed between 60 and 110 mg. The average size of soybeans used was between five to eight millimeters in diameter and spherical. Cotton seeds were more oblong, three to four millimeters in diameter and ten millimeters long. Seeds were surface-sterilized by soaking in 6% bleach solution for 10 minutes in a sterile petri-dish and washed three times with sterile distilled water. Seeds were surface sterilized to maximize the effect of seed exudates on the growth of microbes from the soil. Six replicate seeds were sown into the center of individual wells, halfway into the 15 mm depth of the well, using flamed forceps. Wells containing only soil without a cotton or soybean seed were used as a control. The 12-well microtiter plates were placed in a planting tray covered with a lid to keep the soil from drying. Planting trays containing 12-well microtiter plates were placed inside a growth chamber at 25°C.

### Collection of spermosphere

Spermosphere soil samples were collected at 0, 6, 12, and 18 hours after sowing. Wells containing control soil were sampled as a control, and are hereafter referred to as control soil. Spermosphere soil and control soil samples were collected using an 11 mm cork borer cleaned of soil with 70% ethanol and flame sterilized between samples. The 11 mm cork borer was specifically used since the spermosphere is defined as the first 5-10 mm of soil around a germinating seed (Nelson 2004) and allowed soil collection within this range based on the seed sizes stated previously. Therefore, given the size of the well, the volume of soil used, and the seed sizes, the spermosphere soil sampled consisted of three to six millimeters on either side of a soybean seed and seven to ten millimeters above and below a soybean. Similarly, the spermosphere soil sampled for cotton consisted of seven to eight millimeters on either side and five to ten millimeters above and below the seed. A diagram of the sampling procedure for soybean is shown in Figure 1. In preliminary experiments, bacterial populations in spermosphere soils sampled with this method increased significantly by 1.15 log in soybean and about 0.8 log in cotton compared to control soil (Supplemental Figure 1).

**Figure 1.**
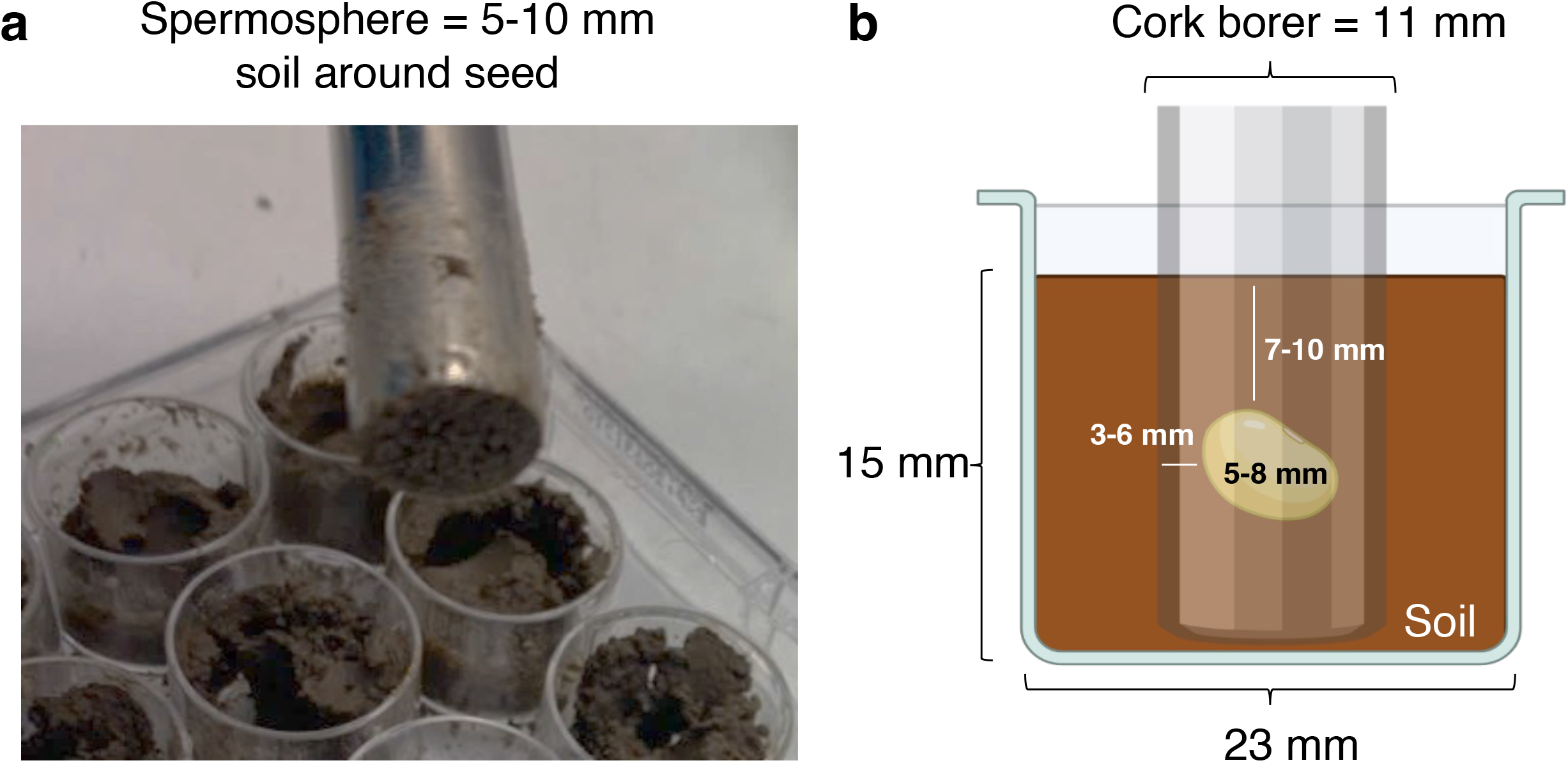
Diagram depicting the sampling technique used for sampling spermosphere soils. The spermosphere is defined as the 5-10 mm of soil directly surrounding a seed. (a) Photo demonstrating the sampling technique of spermosphere soil contained within an 11 mm cork borer. (b) Sampling with an 11 mm cork borer inside the confining space within wells of a 12-well plate allowed direct and controlled sampling of the spermosphere around single seeds.

Spermosphere soil containing the seed inside the core within the cork borer was transferred into sterile envelopes, and 0.25 ml was immediately transferred to 2 ml disruptor tubes (Omega Bio-Tek E.Z.N.A Soil DNA; Norcross GA), then stored at -80ºC until DNA extraction. The remaining soil clinging to the seed was washed off, the seed blotted dry of excess water, and the weight of the seed was recorded after sample collection and compared to the initial individual seed weight to determine the water imbibed by each seed.

### DNA extraction, amplification, and sequencing

The total DNA was extracted from the spermosphere and control soils following the manufacturer’s recommendation. Amplification and library construction of 16S or ITS rDNA was performed with a three-step Polymerase Chain Reaction (PCR) (Lundberg et al. 2013; Longley et al. 2020). Briefly, the 16S region of the ribosomal DNA (rDNA) was amplified using the forward and reverse primers 515F and 806R (Caporaso et al. 2011). Amplification of ITS used the primers ITS1F and ITS4. Following the amplification of the respective rDNA regions, the amplicons were linked to variants of frameshift primers, and then a 10 bp barcode was added for sample identification. Library negative controls consisted of DNA extraction without soil and no template PCR water controls. The ZymoBIOMICS microbial community DNA standard (Zymo Research, Irvine, CA) was used as a positive control mock community. A fungal synthetic mock community was used as a positive control for fungi (Palmer et al. 2018). DNA amplification was confirmed with gel electrophoresis, and successfully amplified libraries were normalized using SequalPrep™ Normalization Plate Kit (Thermo Fisher, USA). Normalized amplicons were then pooled and concentrated 20:1 using the 50K Dalton Millipore filters (Sigma-Aldrich, USA). The pooled library was cleaned using AMPure XP beads at a ratio of 0.7X (Beckman Coulter, USA). Cleaned amplicon pools were verified by gel electrophoresis, quantified using a Qubit fluorometer (Thermo Fisher, USA), and sequenced on an Illumina MiSeq 2×300 bp using the v3 600 cycles kit at SeqCenter LLC (Pittsburgh, PA). Primers and cycling parameters to construct libraries were the same as in Noel (2022).

### Read processing

The quality of demultiplexed reads was assessed using the FastQC, and primer sequences were removed using cutadapt 4.0 (Martin 2011). Prokaryote 16S V4 sequences were merged using VSEARCH 2.21.1 (Rognes et al. 2016). Only forward fungal ITS1 reads were used since reverse reads did not overlap. Fungal reads were trimmed to remove the conserved 18S regions. Reads were then truncated to equal length (fungi 200 bp, prokaryote 300 bp) and quality filtered using VSEARCH 2.21.1 with an expected error threshold of 1.0. Singletons were removed and reads *de novo* clustered based on 97% identity into prokaryote OTUs (pOTUs) or fungal OTUs (fOTUs) using USEARCH v11.0.667, which includes a chimera detection and removal step (Edgar 2010; Edgar et al. 2011). The resulting pOTUs were aligned using MAFFT v7.505 (Nakamura et al. 2018), and a phylogenetic tree was estimated using FastTree v2.1.20 (Price et al. 2010), then midpoint rooted with FigTree v1.4.4 (Rambaut 2018). Taxonomy was assigned to resulting pOTUs using the SINTAX algorithm (Edgar 2016) against the SILVA 138.1 database (Quast et al. 2013). Fungal taxonomy was assigned using the ribosomal database project’s Naïve Bayesian Classifier algorithm against the UNITE fungal ITS database version 9.0 (Nilsson et al. 2019).

### Data analysis

Data were primarily analyzed using phyloseq v.1.34.0 (McMurdie and Holmes 2013) and vegan v2.5-7 (Oksanen et al. 2022) of the statistical software R v.4.2.2. All plots were generated using the data visualization package ‘ggplot2 v.3.3.5’ (Wickham 2016). Contaminant OTUs detected in the negative controls were removed with decontam v1.10.0 (Davis et al. 2018). Samples with less than 10,000 were discarded. Fungal samples with less than 1000 reads were discarded due to low sequencing coverage.

Richness, Pielou’s evenness (Pielou 1966), and Faith’s Phylogenetic Diversity (Faith 1992) were used to determine within sample diversity differences in diversity using Kruskal-Wallis one-way analysis of variance. Read counts were then normalized using the cumulative sum scaling with metagenomeSeq v1.32.0 (Paulson et al. 2013) and subjected to principal coordinate analysis based on Bray-Curtis distances for fungi and prokaryotes or Weighted Unifrac distances for prokaryotes only. This analysis was followed by a Permutational Analysis of Variance (PERMANOVA) implemented with the ‘adonis2’ function to determine the differences in centroids of the prokaryote or fungal communities across time points and soil versus spermosphere. Differences in multivariate dispersion were also evaluated using the ‘betadisper’ function.

Differential abundance analysis was conducted with Analysis of Compositions of Microbiomes with Bias Correction version 2 (ANCOM-BC2) (Lin and Peddada 2020). Significantly different OTUs were detected based on Holm-Bonferonii corrected p-values. Then, microbial co-occurrence networks with prokaryotes and fungi were constructed using SpiecEasi v1.1.2 (Kurtz et al. 2015) and compared between soybean spermosphere soil, cotton spermosphere soils, and control soil using NetCoMi v.1.1.0 (Peschel et al. 2021). For network construction, spermosphere soil samples and soil samples without seeds at 12 and 18 hours were filtered to a common set of taxa with a relative abundance above 0.001% and occupancy above 90%. Co-occurrence association matrices were estimated using the Meinshausen and Bühlmann algorithm with the ‘nlambda’ set to 100, sampled 100 times, and the ‘lambda.min.ratio’ set to 10^−1^. All resulting networks contained stability values of 0.048 or above, close to the 0.05 StARS algorithm stability target. Association matrices for spermosphere soils or control soil were compared using the ‘netAnalyse’ function from NetCoMi. Hub taxa were identified based on eigenvector centrality values above the 95% quantile of a fitted log-normal distribution. Comparison of the hub taxa composition was based on the Jaccard similarity index.

The Data files and scripts used for this analysis are available on GitHub (https://github.com/Noel-Lab-Auburn/SpermosphereMicrobiome2022). Raw sequence reads were deposited to the sequence read archive with the accession number PRJNA925866.

## Results

### Sequencing outputs

Mock OTUs for the fungi and prokaryotes made up 99.9% of the composition of the positive controls, indicating minimal cross-contamination. Nine prokaryotic OTUs were filtered after detection in negative control samples, resulting in 2,090,814 16S V4 reads of 8088 OTUs across 71 samples with a median read depth of 29,237 reads per sample. Nineteen fungal OTUs were detected in negative controls and taken out, resulting in 2,534,301 ITS1 reads with 1904 fungal OTUs across 69 samples and a median read depth of 37,933 reads per sample. Rarefaction curves indicate that much of the diversity was adequately captured (Supplemental Figure 2).

### Prokaryote community dominance correlates with water imbibition

The individual measurement of seed weight for soybean and cotton seeds before and after spermosphere collection indicate that water was imbibed from the surrounding soil (Fig. 2a). Overall, soybean seeds imbibed more (250-300 mg) than cotton seeds (50-80 mg) and both seeds increased in seed weight within the first six hours indicating imbibition within this timeframe, then a plateau after six hours.

**Figure 2.**
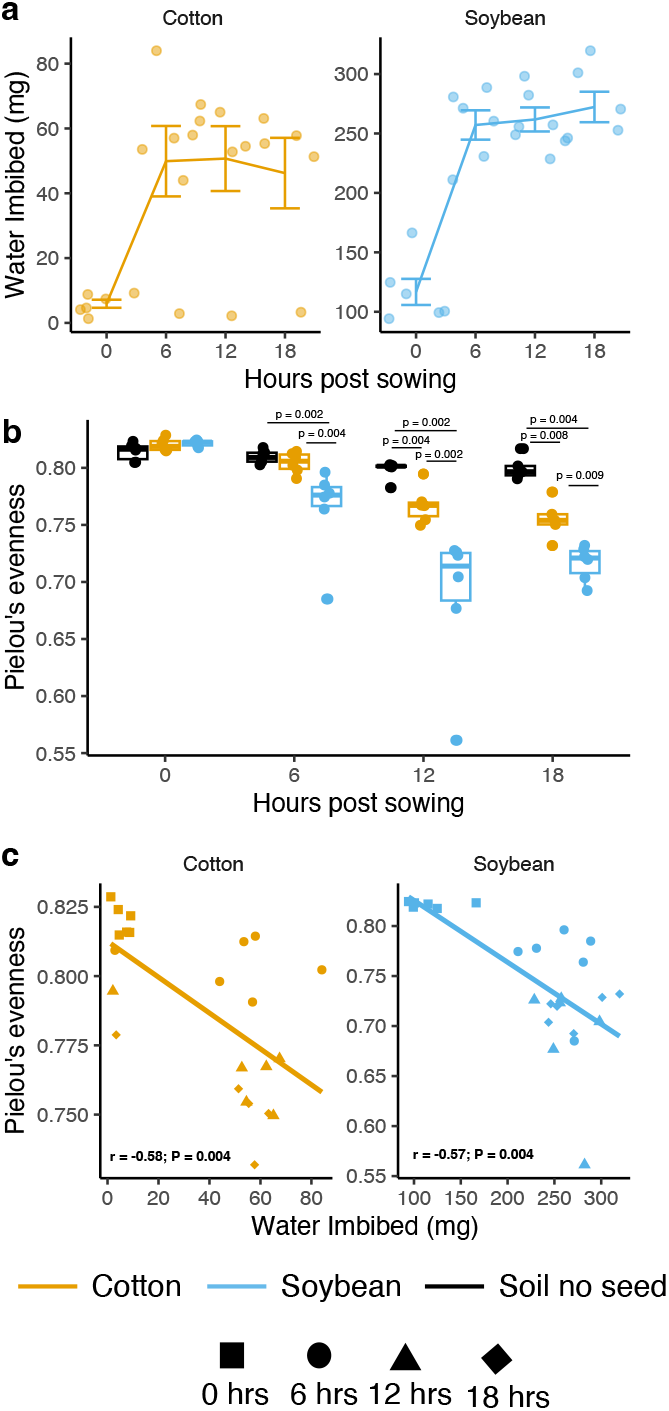
Prokaryote spermosphere evenness follows water imbibition. (a) Water imbibition over time for cotton and soybean seeds (n = 6). (b) Prokaryote evenness over time in control soil, soybean spermosphere soil, or cotton spermosphere soil. Soybean had significantly lower evenness (higher dominance) after 6, 12, and 18 hours compared to control soil. Cotton followed the same trend but was significantly less even after 12 and 18 hours. Comparisons were made with Wilcox ranked sign test (α = 0.05; n = 6) (c) Spearman correlation between prokaryote evenness and water imbibition.

We observed a reduction in prokaryote evenness (or increase in dominance) over time in spermosphere soils but not in control soil (Fig. 2b). At time-point 0, there was no significant difference in the evenness of prokaryote taxa (*P* = 0.18), as expected. At 6 hours, we observed a reduction in the evenness of prokaryote taxa in soybean spermosphere soils, compared to cotton spermosphere (*P* = 0.004) or control soil (*P* = 0.002). At 12 and 18 hours, both cotton and soybean spermosphere samples had significantly reduced evenness compared to control soil and each other (*P* ≤ 0.009). Prokaryote evenness was significantly negatively correlated with water imbibition, meaning that as seeds imbibed water and released exudates, prokaryote communities became more dominant (cotton r = -0.58, *P* < 0.001; soybean r = -0.57, *P* = 0.004) (Fig. 2c). However, the crop did not alter prokaryote richness or phylogenetic diversity compared to control soil. Prokaryote richness and phylogenetic diversity dropped significantly over time regardless of habitat (*P* < 0.001). Additionally, there was no consistent evidence that fungal richness or evenness was altered in spermosphere samples compared to control soil. Still, a few samples of soybean spermosphere soil and control soil dropped in evenness after 18 hours (Supplemental Figure 3), becoming more dominant in one fOTU2 *Fusarium* (Supplemental Figure 4).

### Spermosphere prokaryote composition depends on the crop

Spermosphere soils had different prokaryote community compositions than control soil. A visualization of the change in the most abundant prokaryote composition over time is shown in Figure 3. Prokaryote communities were driven by habitat (*P* < 0.001), time since sowing (P < 0.001), and interaction of both factors (P < 0.001) (Supplemental Table 1). The interaction prompted a closer look into the differences observed between crops by splitting the data by time-point (Figure 4; Table 1). At 0 hours, no significant difference in prokaryote communities existed between bulk soil compared to soybean and cotton spermosphere, as expected (Bray-Curtis, *P* = 0.452; Weighted Unifrac, *P* = 0.192). However, as early as 6 hours, we observed significant differences between control soil and spermosphere soil samples (Bray-Curtis, *P* < 0.001; Weighted Unifrac *P* < 0.001). Differences were extended through 12 hours (Bray-Curtis, *P* < 0.001; Weighted Unifrac *P* < 0.001) and 18 hours (Bray-Curtis, *P* < 0.001; Weighted Unifrac *P* < 0.001), where it was clear the spermosphere formed unique community compositions within soybean or cotton. Further, differences in multivariate dispersions were not observed supporting true differences in centroids rather than group dispersions (Table 1). This same trend was not observed with fungi. Time did alter fungal community composition (*P* = 0.01), but there was no evidence that soybean or cotton altered fungal community composition compared to control soils (*P* = 0.09).

**Table 1.** Permutational Analysis of Variance (PERMANOVA) and dispersion for prokaryote communities separated by time post sowing based on Bray-Curtis and weighted Uni-frac distances

| Dissimilarity<br>Metric | Hours post<br>sowing | PERMANOVA<sup>a</sup> |  |  |  | Dispersion |  |
| --- | --- | --- | --- | --- | --- | --- | --- |
|  |  | df | R<sup>2</sup> | F | P | F | P |
| Bray-Curtis | 0 | 2 | 0.113 | 0.959 | 0.437 | 0.937 | 0.407 |
|  | 6 | 2 | 0.273 | 2.815 | <0.001 | 0.539 | 0.606 |
|  | 12 | 2 | 0.424 | 5.517 | <0.001 | 0.828 | 0.533 |
|  | 18 | 2 | 0.417 | 4.648 | <0.001 | 0.092 | 0.911 |
| Weighted<br>Unifrac | 0 | 2 | 0.142 | 1.238 | 0.192 | 1.898 | 0.190 |
|  | 6 | 2 | 0.491 | 7.245 | <0.001 | 2.538 | 0.066 |
|  | 12 | 2 | 0.609 | 11.675 | <0.001 | 1.186 | 0.376 |
|  | 18 | 2 | 0.696 | 14.863 | <0.001 | 1.401 | 0.304 |

<sup>a</sup>Permanova and dispersion tests were performed on 999 permutations

**Figure 3.**
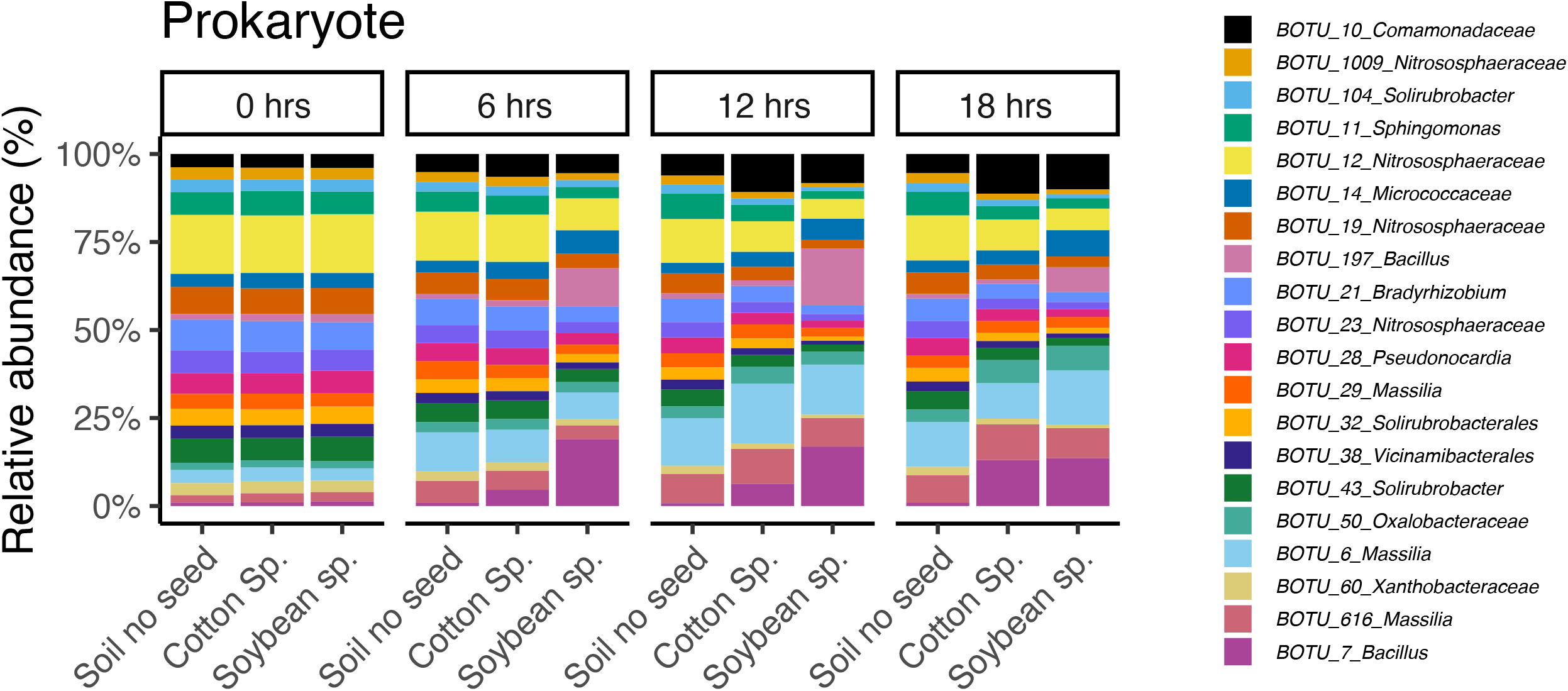
Composition of the most abundant prokaryote OTUs changes over time. Relative abundance of the top twenty most abundant prokaryote OTUs shifts over time within soybean spermosphere soil, cotton spermosphere soil, or control soil.

**Figure 4.**
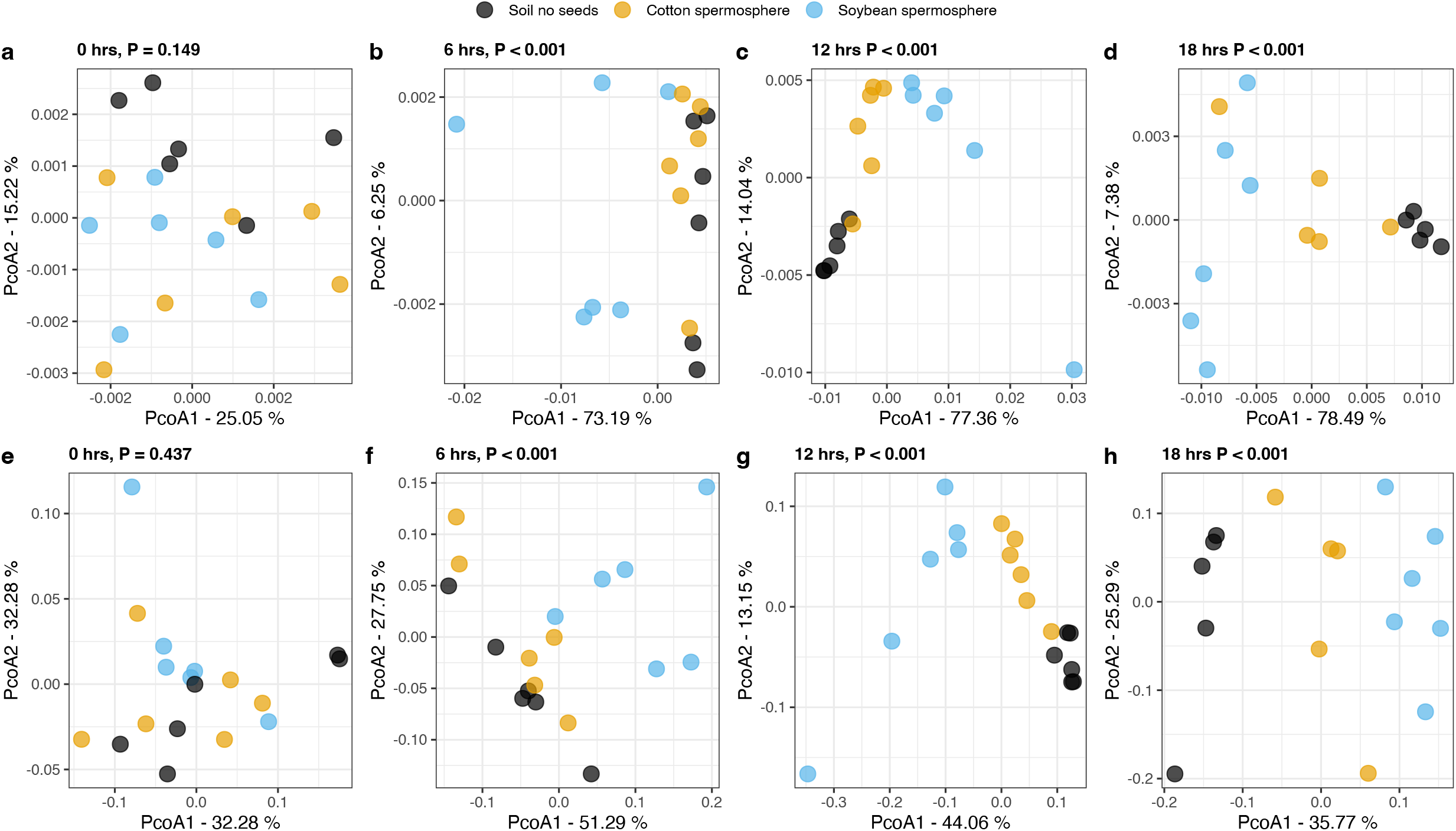
Prokaryote spermosphere composition changes over time and crop. (a-d) Principal coordinate analysis based on Weighted Unifrac distances. (e-h) Principal coordinate analysis based on Bray-Curtis distances. Reported significance values above each plot are the result of a permutational test of variance (α = 0.05; n = 6). Accompanying dispersion tests are shown in Table 1.

### Enriched prokaryotes in the spermosphere have unique and shared taxa among crops

Differential abundance analysis determined sets of pOTUs significantly enriched in the spermosphere of cotton and soybean compared to control soil (Figure 5a; Supplemental Table 2. Ninety-four percent of the enriched taxa belonged to Proteobacteria (57%, n = 27) and Firmicutes (36%, n = 17). The remaining three belonged to the Actinobacteria. Within the Proteobacteria, the enriched taxa were spread across ten prokaryote families, with the most enriched taxa in the Oxalobacteraceae (41%, n = 11). The majority of the enriched Proteobacteria were unidentified at the genus level (n = 12) but included *Massilia* (n = 3), *Noviherbaspirillum* (n = 2), *Burkholderia*/*Paraburkholdaria* (n = 2), *Aquabacterium* (n = 1), *Pseudomonas* (n = 2), *Cupriavidus* (n = 1), *Pantoea* (n = 1), *Paucimonas* (n = 1), *Rubellimicrobium* (n = 1), and *Azospirillum* (n = 1) (Figure 5b). Within the Firmicutes, all but one pOTU belonged to the Bacilli class with the genera *Paenibacillus* (n = 8), *Bacillus* (n = 4), *Brevibacillus* (n = 1), *Exiguobacterium* (n = 1), and *Tumebacillus* (n = 1). Many enriched taxa were shared between cotton and soybean (n = 18), indicating that similar taxa take advantage of releasing exudates from seeds (Figure 5b). All the enriched pOTUs were present in control soil samples meaning it was unlikely they originated from the seed but were present in the soil and proliferated upon exudate release from the seeds.

**Figure 5.**
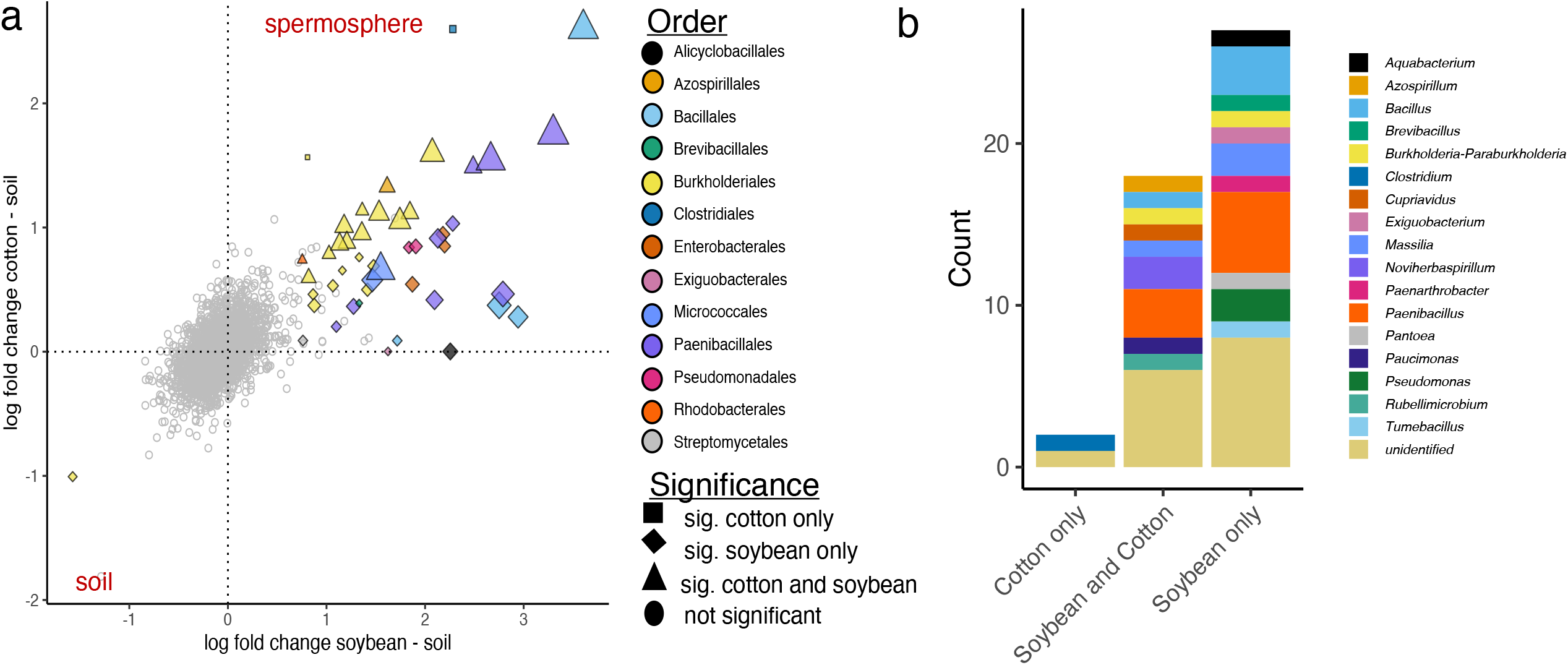
Differential abundance of prokaryote OTUs shows enrichment of specific taxa within the spermosphere. (a) Points represent individual OTUS. Positive values on the x-axis indicate the OTU was numerically more abundant in a soybean spermosphere compared to soil without a soybean seed. Similarly, positive y-axis values indicate the OTU was numerically more abundant in a cotton spermosphere compared to soil without a cotton seed. Colored points are pOTUs detected as significantly different in soybean or cotton. Grey circle points are non-significant. Point shape indicates significance in one crop or both. Points are colored by the prokaryote order. Significance was determined within the ANCOM-BC2 algorithm with a Holm-Bonferroni correction (α-0.05). (b) Composition of significantly enriched pOTUs colored by genera.

### Cotton and soybean spermosphere networks are more connected and have distinct microbial hub taxa

Cotton and soybean spermosphere networks were compared to each other and to the control soil to determine if they contained different topologies, different sets of network hubs, and the centrality of spermosphere-enriched taxa. Overall, network topology parameters were similar between networks except for the number of separate components. In other words, the spermosphere soil networks mainly consisted of one more prominent component and fewer disconnected sub-networks than the control soil network (Figure 6). For example, the control soil network contained 30 components and 80 nodes within the largest component. Soybean and cotton spermosphere soil networks had more nodes within the largest component (cotton = 136, soybean = 121). However, control soil and a slightly higher positive edge percentage (61% without a seed, 58% cotton spermosphere, 52% soybean spermosphere) (Supplemental Table 3).

**Figure 6.**
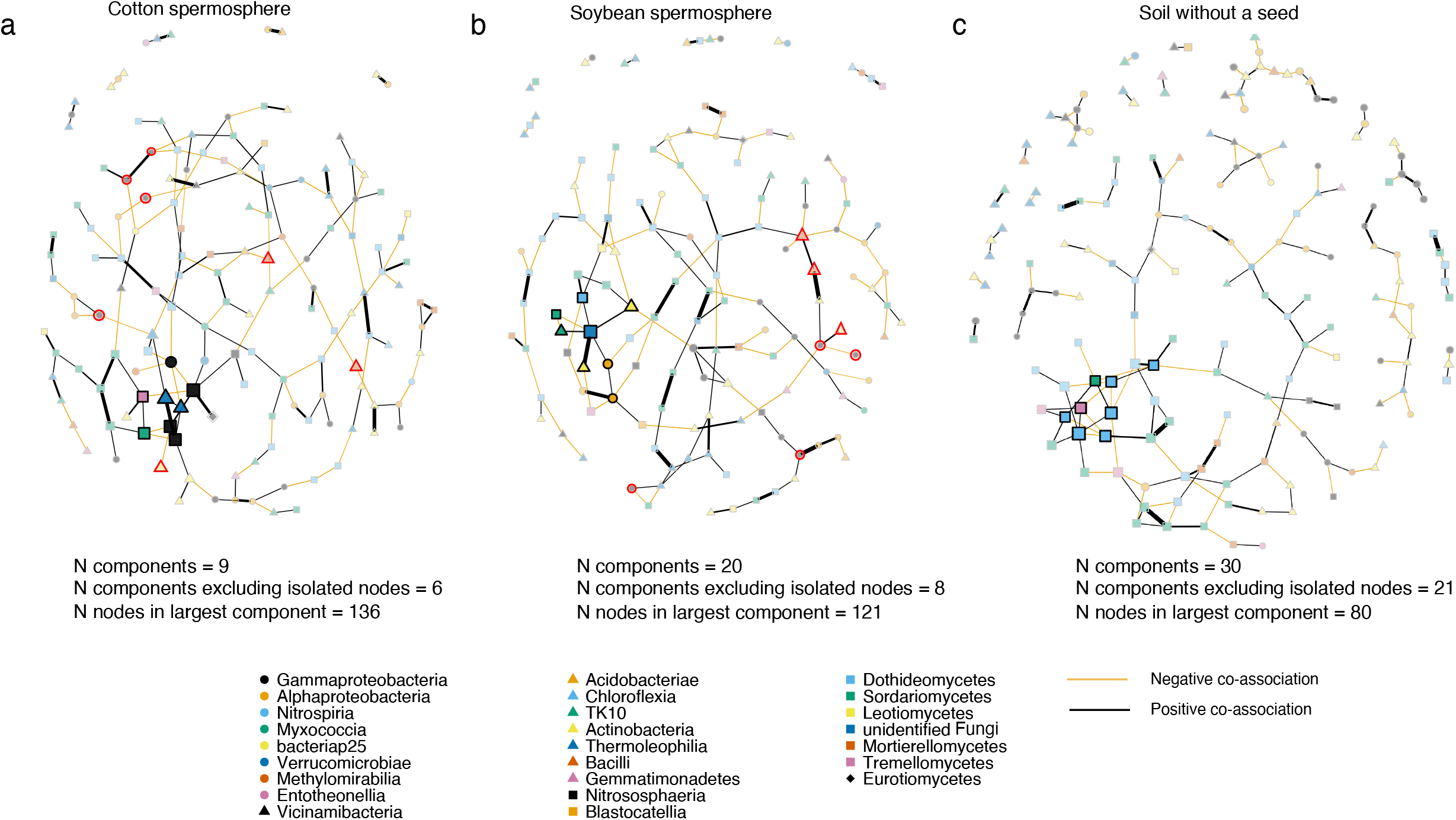
Spermosphere soils have different topological properties and different sets of hub taxa compared to control soil. Networks for (a) cotton spermosphere soil and (b) soybean spermosphere soil and (c) control soil were constructed with the same set of taxa. Nodes with different shape and colors indicate the prokaryote or fungal class. Less transparent nodes are significant hubs based on eigenvector centrality above the 95% quantile. Nodes with red outlines were significantly enriched in a spermosphere as detected in Fig.4.

Comparison between central nodes was significantly different, indicating that the hub taxa were different between networks given the same sets of taxa used to construct the networks (Table 2). Hub taxa for cotton consisted of six prokaryote OTUs and two fungal OTUs. Cotton prokaryote hubs consisted of three Archaea OTUs in the Nitrososphaeraceae family (pOTU1009, pOTU19, and pOTU12), two OTUs from the Gaiellales (pOTU46, pOTU119), and pOTU29 *Massilia* (Figure 6a; Table 3). Fungal cotton hubs were fOTU56 *Fusarium chlamydosporum*, and a yeast fOTU36 *Hannaella oryzae* (Figure 6a; Table 3), whereas the soybean network had three fungal hub taxa: fOTU64 *Helicoma*, fOTU10 *Bartalinia pondoensis*, and an unidentified Fungus fOTU36 (Figure 6b; Table 3). Prokaryote hub taxa in the soybean spermosphere network contained pOTU11 *Sphingomonas*, pOTU132 Nocardioides, pOTU1559 Chloroflexi TK10, pOTU36 Angustibacter, and pOTU349 *Methylobacterium*/*Methylorubrum* (Figure 6b; Table 3). The network from the control soil contained only fungal hubs, different than the identities of spermosphere fungal hubs except for fOTU36 *Hannaella oryzae*. Spermosphere-enriched taxa included in the network analysis were not hub taxa indicating that although enriched in a spermosphere, other microbial taxa besides the enriched taxa play an essential role in maintaining spermosphere network structure (Figure 6).

**Table 2.**
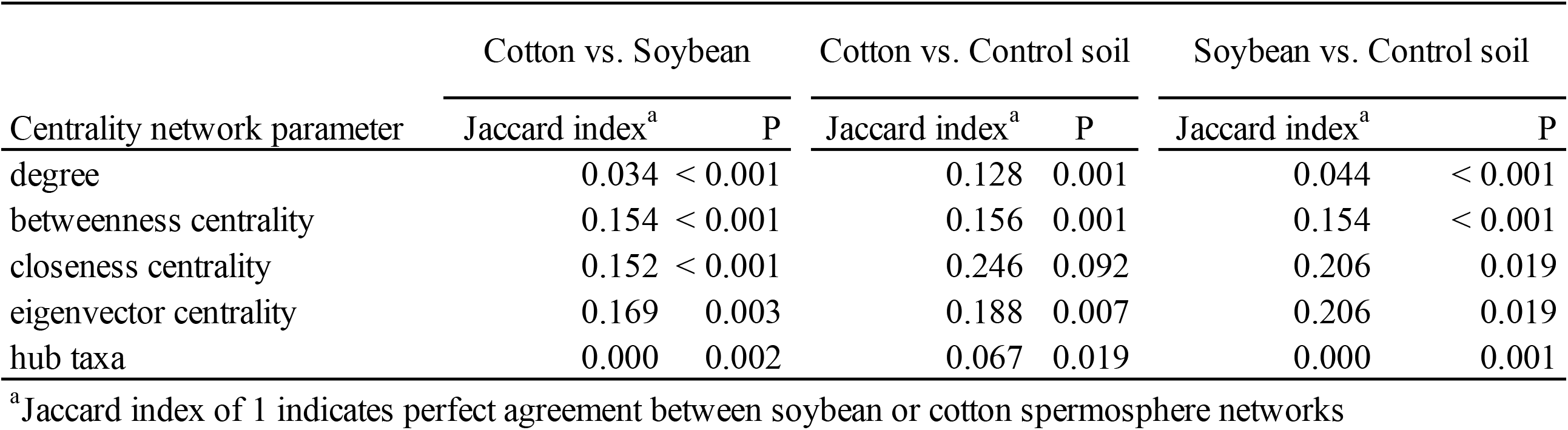
Comparison of network centrality parameters

**Table 3.** Hub taxa identified for each habitat

| Crop | OTU | Taxonomy |
| --- | --- | --- |
| Cotton<br>spermosphere soil | pOTU46 | Gaiellales |
|  | pOTU119 | Gaiellales |
|  | pOTU29 | *Massilia* |
|  | pOTU1009 | Nitrososphaeraceae |
|  | pOTU19 | Nitrososphaeraceae |
|  | pOTU12 | Nitrososphaeraceae |
|  | fOTU36 | *Hannaella oryzae* |
|  | fOTU56 | *Fusarium chlamydosporum* |
| Soybean<br>spermsphere soil | pOTU1559 | Chloroflexi TK10 |
|  | pOTU132 | Nocardioides |
|  | pOTU36 | Angustibacter |
|  | pOTU11 | *Sphingomonas* |
|  | pOTU349 | *Methylobacterium /Methylorubrum* |
|  | fOTU64 | *Helicoma* |
|  | fOTU36 | Fungi |
|  | fOTU10 | *Bartalinia pondoensis* |
| Soil without a seed | fOTU1 | *Stagonosporopsis oculi-hominis* |
|  | fOTU13 | Teichosporaceae |
|  | fOTU15 | Cucurbitariaceae |
|  | fOTU17 | *Alternaria tenuissima* |
|  | fOTU22 | Neopestalotiopsis |
|  | fOTU32 | Pleosporales |
|  | fOTU36 | *Hannaella oryzae* |
|  | fOTU4 | *Cladosporium cladosporioides* |

## Discussion

To our knowledge, this is the first study to use culture-independent sequencing to study soybean and cotton spermosphere soil microbiomes during the first phases of seed germination. The advancement that allowed this was the method that constrained non-sterile soil to wells within a 12-well plate and sampled around an imbibing seed with a cork borer. The technique enabled the precise and efficient collection of spermosphere soils as defined in space and time (Nelson 2004), which we believe this represents a more realistic spermosphere habitat. The focus on spermosphere soil in the first phases of seed germination differs from other studies that pre-imbibe or pre-germinate seeds in axenic conditions. We hypothesized and observed a rapid increase in water imbibition followed by a plateau characterizing phase I and phase II of seed germination. Prokaryote community structure changed in as little as six hours for soybean and twelve hours for cotton. We did observe that crops had unique prokaryote community structures in the spermosphere that were distinct from the control soil, typified by differences in network hub taxa and network topologies. The differing hub taxa demonstrate that others besides the enriched taxa are integral to each crop’s spermosphere community structure. However, despite the differences in composition and hub taxa, among the most important observations was the commonality in the enriched copiotrophic taxa with a long history of benefiting plant growth, such as *Bacillus, Paenibacillus, Burkholderia, Massilia, Azospirillum*, and *Pseudomonas*.

In this study, we further define the development of the spermosphere of cotton and soybean at six to twelve hours after sowing, which aligns with previous studies of increased spore germination and full colonization of cotton seeds by *Pythium ultimum* twelve hours after sowing (Nelson 1986, 1988). We observed an increase in water imbibed by both cotton and soybean seeds in the first six hours, which is consistent with previous reports that documented increased water imbibition and exudation within the first few hours after sowing (Simon and Raja Harun 1972). Imbibition ceased representing phase II of germination, indicating the saturation of nutrient reserves and synthesis of products required for the extending radicle (Nelson 2004).

Similar to several other studies, we observed the soil microbes respond to seed exudates and dominated the spermosphere microbiomes (Buyer et al. 1999; Hardoim et al. 2012; Ofek et al. 2011). We observed changes in phylogenetic dissimilarity between crops, and since phylogenetically similar species are more likely to share ecological characteristics and functional traits (Morrissey et al. 2016), it may be expected that the spermosphere communities in our study changed in a functional capacity as in Buyer et al. (1999). However, we observed varying spermosphere prokaryote composition in different plant species, which may highlight the importance of sample collection at the initial stages of seed germination and imbibition rather than at later hours potentially after radicle emergence. Additionally, as in Buyer et al. (1999), future studies should also include other soils with inherently different communities to understand the contribution of different soil microbial pools to forming the spermosphere.

The difference between crops may have also been due to differences in the amount of water imbibed. We noticed that soybean imbibed more than cotton seeds, likely due to seed size (Soldan et al. 2021; Vančura 1972). Vančura and Hanzlíková (1972) demonstrated increased quantities of seed exudates as seed size increased. Different varieties of common bean have been shown to differ in the amount of seed exudates, with larger seeded varieties releasing more exudates (Kato et al. 1997). Thus, we speculate that the greater and faster turnover in microbial communities of the soybean spermosphere compared to cotton may be due to the larger size of soybean seeds and increased exudation, which potentially supported a larger habitat for the microbes to occupy. It also leaves the question if microbial communities would have converged on similar compositions if a later sampling point was included.

Regardless, as a result of water imbibition and seed exudation, we observed a change in dominance in the spermosphere microbiome over time with both crops. Ota et al. (1991) showed specific nitrogen-fixing bacteria had increased dominance in the spermosphere of cocklebur seeds but not in soil. Upon revealing enriched taxa in soybean and cotton spermosphere soils, we found some commonalities. Importantly, Bacilli was enriched in both crop’s spermosphere soils. Since these *Bacilli*, including *Tumebacillus, Paenibacillus*, and *Bacillus*, have historically been associated with plant growth promotion and disease protection and have commercial potential, they were notable. It indicates their ability to utilize seed exudates quickly for growth. Seed exudates have been reported to induce chemotaxis, seed colonization, and biofilm formation of *B. amyloliquefaciens* (*velezensis*) by enhancing active cell division (Yaryura et al. 2008). *Paenibacillus polymyxa* isolated from wheat and peanut rhizosphere increased the survival of *Arabidopsis thaliana* in the presence of the oomycete pathogen *Pythium aphanidermatum* when applied as root treatment (Timmusk et al. 2009). Identifying these enriched taxa is important for prioritizing future work on a mechanistic understanding of the spermosphere microbial interactions that will improve the development of efficacious biologically based disease solutions (Nelson 2004; Weller 2007).

In terms of seed versus soil origin, there were OTUs with a low relative abundance and low occupancy that only occurred in cotton or soybean spermosphere samples and were absent from the soil. However, we hesitate to conclude they originated from the seed without directly identifying seed epiphytes and endophytes since it was impossible to know if the unique microbes were seed associates colonizing the spermosphere or if rare members of the soil only present spermosphere samples by chance. Furthermore, surface disinfecting seeds used in this study likely reduced the number of seed epiphytes that would colonize the spermosphere. The implications of surface disinfecting seeds have been argued elsewhere (Barret et al. 2015; Schiltz et al. 2015; Shade et al. 2017). Another limitation of our approach that limited our ability to identify seed-associated microbes may be the use of OTUs rather than amplicon sequence variants (ASVs) or zero radius OTUs (zOTUs). A finer clustering method may be better suited to studying the transmission of seed-associated microbes into the spermosphere since genotypes originating from the seed may be grouped within 97% OTUs originating from the soil. We recognize microbes originating from the seed can colonize seedlings and other plant organs, which can alter plant health (Bintarti et al. 2022b; Bintarti et al. 2022a; Chesneau et al. 2022; Chesneau et al. 2020; Johnston-Monje et al. 2016; Mitter et al. 2017; Rochefort et al. 2021; Shade et al. 2017; Simonin et al. 2022). For example, it was recently demonstrated that crop flowers sprayed with a beneficial bacterium can colonize endosperm and transmit to germinating seeds (Mitter et al. 2017). While the importance of seed-associated microbes on plant health is evident, little is known about seed endophytes and interactions with horizontally acquired soil organisms, which tend to contribute a large portion of the microbial diversity to the seedling microbiome (Buyer et al. 1999; Nelson 2018; Rochefort et al. 2021).

In terms of the microbial networks, we observed different hubs and different topologies given the same set of taxa used for network construction. While fungal diversity was not altered in this study, fungal OTUs were identified as hubs, potentially demonstrating meaningful interactions with in a spermosphere soil. Of most interest was the yeast *Hannella* since these fungi are commonly found in soils, the phyllosphere, and as part of the core seed and phyllosphere microbiome (Noel et al. 2022; Simonin et al. 2022; Yurkov 2018). *Dioszegia*, in the same family as *Hannaella*, was identified as a network hub in the phyllosphere (Agler et al. 2016), and the closely related yeast *Bullera* has been a network hub of the soybean phyllosphere (Longley et al. 2020). These yeasts are generally non-pathogenic, but their ecological role is poorly understood (Gouka et al. 2022). The prokaryote hubs were also intriguing because cotton contained several Nitrososphaeraceae pOTUs, which likely are involved with ammonia-oxidizdation in soils (Reyes et al. 2020). Cotton spermosphere hubs also had a *Massilia* pOTU. *Massilia* is known for below-ground associations and the ability to solubilize phosphate (Silva et al. 2017), but has also been found as a hub in above-ground plant tissues (Longley et al. 2020). For soybean, *Sphingomonas* and *Methylobacterium*/*Methylorubrum* pOTUs were notable network hubs since these genera have been demonstrated to be abundant in the phyllosphere and core seed microbiome and produce plant growth-promoting hormones and UVA-absorbing compounds (Kwak et al. 2014; Yoshida et al. 2017). The difference in hub taxa between crops demonstrates that soybean and cotton construct unique microbial communities early in life, which may have plant health consequences at or beyond the spermosphere stage.

However, spermosphere-enriched pOTUs were not identified as network hubs; instead, they were located more peripheral in the networks, indicating they may be copiotrophs responding quickly to the availability of carbon-rich exudates from the seeds (Torres-Cortes et al. 2018). Spermosphere networks were more connected with larger components than the soil network. Increased soil network complexity was associated with increased microbiome function (Wagg et al. 2019). Therefore, it may be hypothesized that seed exudates help stimulate associations between organisms or sub-communities and form more connected or stable communities. However, further research is needed to determine how topological features of networks are associated with plant health and why hub taxa connect to other taxa and help assemble plant microbiomes.

The technique used in this study enabled quick and efficient collection of spermosphere soil within phase I and II of seed germination and showed the enrichment of beneficial copiotrophic taxa. However, these copiotrophic taxa were not central to microbial networks. This technique could easily be applied to other sequencing methods like metagenomics or metatranscriptomics for a better understanding of spermosphere soil microbiome functions. Coupled with sequencing the seed microbiomes will be powerful to study interactions between seed and seedling pathogens, chemical or biological seed treatments, and interactions with pathogens in the spermosphere – thereby improving knowledge of spermosphere ecology, which will lead to improved understanding of the plant microbiome.

## Acknowledgement

This work is/was supported by the USDA National Institute of Food and Agriculture, Hatch project 1025628. Additional support comes from the Alabama Farmer Federation Soybean and Cotton Committees and the Department of Entomology and Plant Pathology at Auburn University.

## Table and Figures

**Supplemental Table 1**. Permutational Analysis of Variance (PERMANOVA) for prkaryote and fungal communities

**Supplemental Table 2**. Differentially abundant bacterial taxa in soybean and cotton spermosphere soils

**Supplemental Table 3**. Topological properties of networks

**Supplemental Figure 1**. Preliminary experiment conducted to demonstrate the effectiveness of the sampling procedure shown in Figure 1. Bacterial populations within spermosphere were greater in soybean or cotton spermosphere soils compared to control soil.

**Supplemental Figure 2**. Sequencing outputs for prokaryotes (16S) and fungi (ITS). (a) Composition and taxonomic output of the mock community samples. (b) rarefaction curves for prokaryotes and fungi. Dashed lines indicate the median read depth. (c) contaminant filtering based on OTU prevalence in negative control samples. (d) Histogram of read depth per sample.

**Supplemental Figure 3**. Within sample diversity measurements for soybean or cotton spermosphere samples compared to control soil. (a) prokaryote richness significantly dropped over time (P < 0.001) consistently across habitats (spermosphere or soil without seed). (b) Similarly, Faith’s phylogenetic distance also followed a similar pattern. No consistent differences were observed over time or between habitats for fungal (c) richness or (d) evenness.

**Supplemental Figure 4**. Fungal composition of the top 20 most abundant Fungal OTUs in cotton spermosphere soil, soybean spermosphere soil, or control soil.

