## Supplementary figures and images for "Soybean and cotton spermosphere soil microbiome shows dominance of soil-borne copiotrophs"

### Supplemental Figure 1

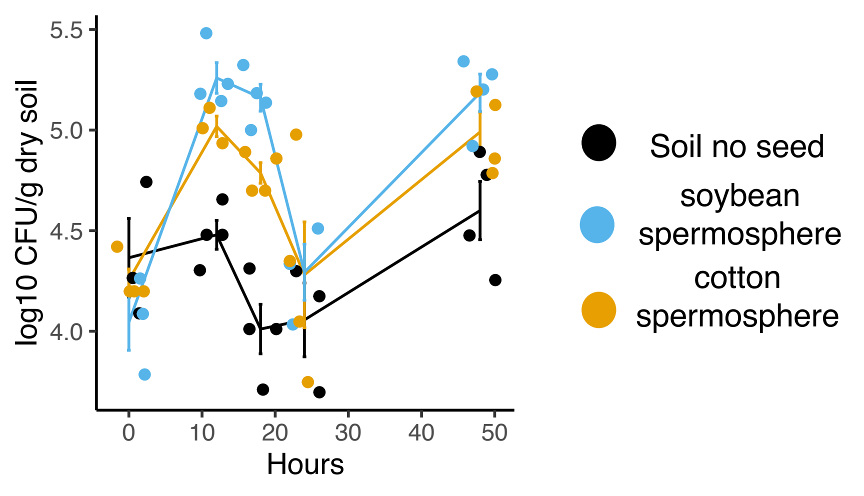

### Supplemental Figure 2

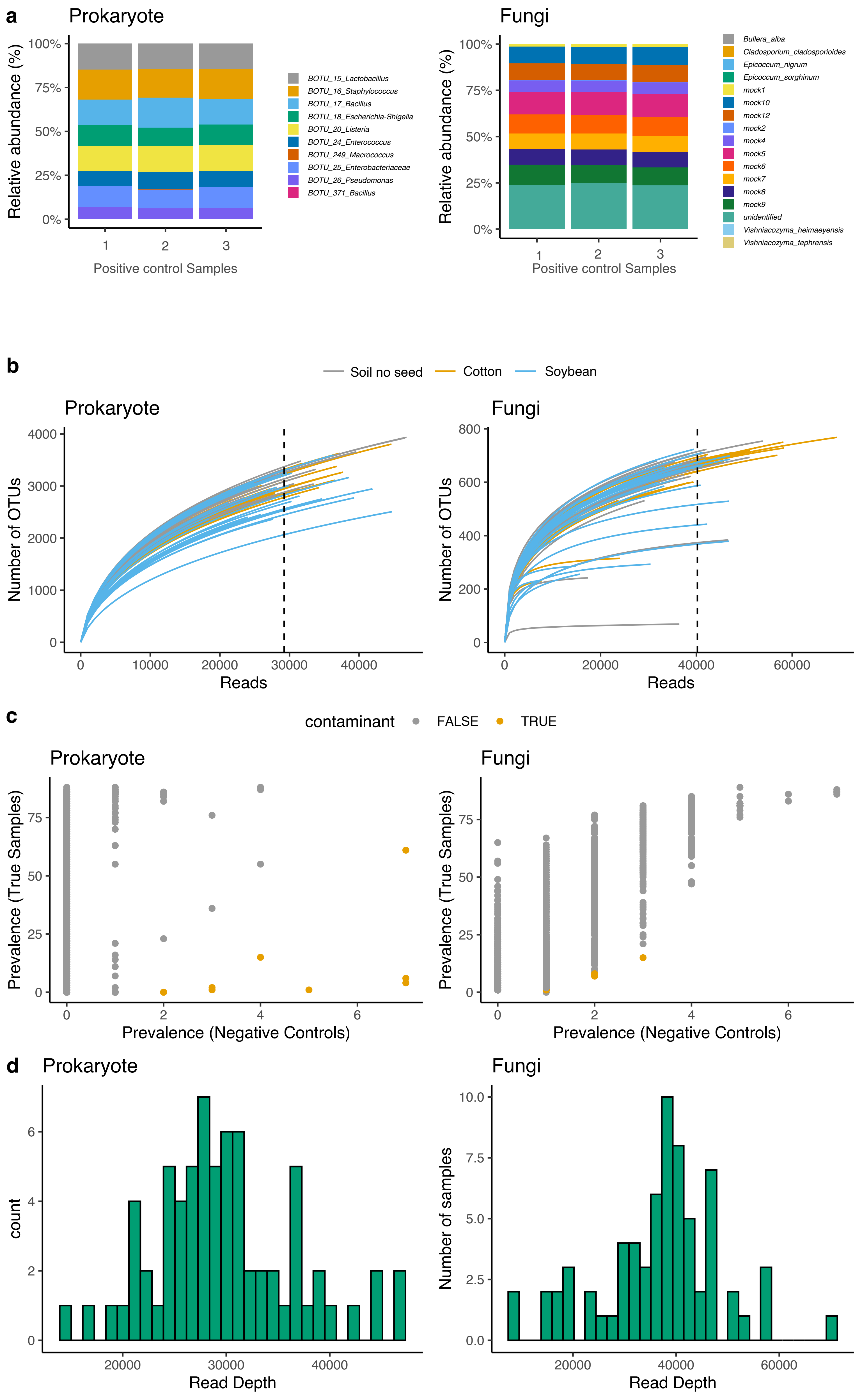

### Supplemental Figure 3

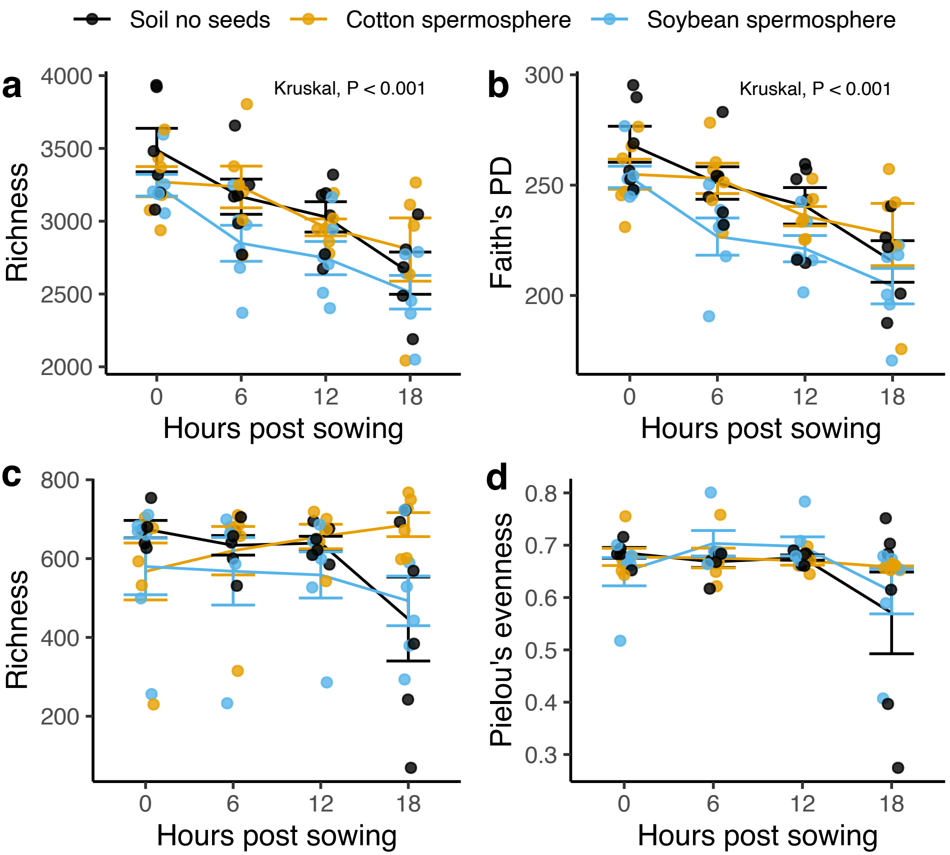

### Supplemental Figure 4

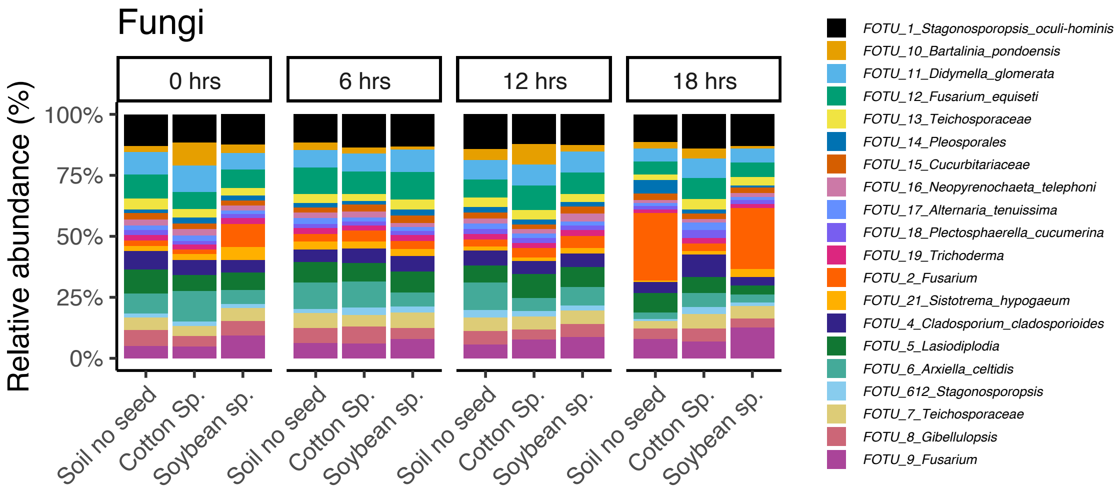
